# Targeting Innate Immune Signaling in Glioma-Initiating Cells Impairs Self-Renewal and Radiation-Induced Cellular Plasticity

**DOI:** 10.1101/2024.12.13.628405

**Authors:** Anjelica M. Cardenas, Linda Azizi, Angeliki Ioannidis, Kruttika Bhat, Ling He, Mahya Mohammadi, Frank Pajonk

## Abstract

Organisms constantly face environmental stressors that threaten their cellular and genomic integrity. In their response, pathogen-associated molecular patterns (PAMPs) and/or damage-associated molecular patterns (DAMPs) are detected by pattern recognition receptors (PRRs) and trigger the innate immune response. In this study we tested the hypothesis that DAMPs contribute to radiation-induced cellular plasticity in Glioblastoma (GBM). GBM is known to be organized hierarchically with a small number of glioma-initiating cells (GICs) driving treatment resistance and recurrences.

Using patient-derived GBM specimens, we employed sphere forming capacity assays and *in vitro* extreme limiting dilution assays to examine how innate immune receptor signaling impacts the maintenance and self-renewal of GICs. By leveraging an imaging system for putative GICs we determined *de novo* induction of GICs from non-stem glioma cells.

We find that GIC maintenance after irradiation is mediated by cGAS-independent STING signaling, possibly involving signaling through TLR4 and TLR9. Induction of radiation-induced plasticity involves TLR3 signaling, with potential roles for other receptors and processes modulated by MyD88. These findings suggest that targeting innate immune signaling could prevent radiation-induced cellular plasticity for potential therapeutic benefit.

## Introduction

Life evolved in the presence of constant genomic injuries. To survive, normal cells and tissues have developed effective strategies to repair DNA damage and replace dying cells to maintain tissue integrity. In tissues organized hierarchically, the stem cell compartment replaces dying cells with newly differentiating cells. But even if the stem cell compartment is compromised, cellular plasticity allows differentiating cells to regain stem cell traits and replace dying stem cells. When essential developmental transcription factors, termed *Yamanaka Factors* (YFs) are re-expressed, even terminally differentiated, somatic cells regain pluripotency [1]. Yet, elegant studies by Cooke *et al*. demonstrated that the re-expression of Yamanaka factors alone is insufficient for reprogramming and requires the presence of danger signals in order for differentiated cells to regain stem cell traits [2]. These danger signals are classified as pathogen-associated molecular pattern (PAMPS) and damage-associated molecular pattern (DAMPS) and signal through pattern recognition receptors (PRRs) of the innate immune system, leading to the production of reactive oxygen species (ROS). Inhibiting the production of ROS impairs the efficacy of nuclear reprograming [3].

Glioblastoma (GBM) is the deadliest adult brain cancer with unacceptable low median survival rates [4]. The current standard of care for GBM involves surgical tumor resection followed by combination treatment of temozolomide (TMZ) and irradiation. Intratumoral heterogeneity, the highly infiltrative nature of GBM, and the presence of treatment-resistant glioma-initiating cells (GICs) contribute to limited treatment success [5]. Given the low median survival rates there is an urgent need to understand the mechanisms driving the aggressive nature of GBM and enable the development of more effective, targeted treatments.

Cellular plasticity and reprogramming are crucial to cancer as they allow tumors to adapt [6]. We previously reported that ionizing radiation (IR) promotes the conversion of non-tumor-initiating cells (non-TICs) into tumor-initiating cells (TICs) in breast cancer and GBM [7, 8]. IR is a form of genotoxic stress that releases DAMPs and activates inflammatory signaling pathways [9].

We hypothesized that DAMPs released after IR activate conserved innate immune signaling pathways that promote cellular plasticity and multipotency. To investigate this, we examined the effects of receptor inhibition on stemness in patient-derived GBM cell lines using sphere formation assays, limiting dilution assays, and reprogramming assays. Results from our study indicate an involvement of the innate immune receptors in modulating GBM cell plasticity in response to irradiation.

## Material and Methods

### Cell lines

Cell lines used (HK-374/345/308/157) were a gift from Dr. Harley I. Kornblum and previously characterized [10]. For monolayers, cells were grown in DMEM media (Gibco, Waltham, MA, USA) supplemented with 10% fetal bovine serum (Seradigm, Avantor, Inc., Radnor, PA, USA) and 1% pen/strep (Gibco) solution, and grown in a humidified 37°C incubator with 5% CO_2_. During passaging, 0.25% Trypsin-EDTA (1x, Gibco) was used to detach the cells. Glioma spheres were grown under similar incubation conditions, in DMEM/F-12 (1:1) (Gibco) media supplemented with 1% pen/strep, 20 ng/ml fibroblast growth factor (bFGF, Sigma, Burlington, MA, USA), 20 ng/ml epidermal growth factor (EGF, Sigma), one bottle of SM1 neuronal supplement containing vitamin A (StemCell Technologies, Vancouver, Canada) and heparin. For passaging spheres, TrypLE^TM^ Express (Gibco) was used. All cell lines were routinely tested for mycoplasma using mycoplasma specific PCR testing and/or MycoAlert (Lonza, Walkersville, MD, USA). Cell line identity was confirmed by DNA fingerprinting (Laragen, Culver City, CA).

### Drug treatments

Information regarding the manufacturers and catalog numbers of the inhibitors used can be found in **Supplementary Table 1**. TLR4i arrived as 10mM stock solution in DMSO, all other drugs were prepared as 10 mM stock solutions in DMSO (TLR3i, CQ, cGASi and STINGi) or HyClone water (TLR9i). Control cells were treated with either DMSO or HyClone water. For TLR3 and TLR4 inhibitors, cells were treated for five consecutive days starting one day after plating since TLR3i and TLR4i had minimal effects when given as a single treatment, likely due to the short half-life of these drugs. CQ was administered once, 24 hours after plating.

### Irradiation

Cells were irradiated at room temperature using an experimental X-ray irradiator (Gulmay Medical Inc. Atlanta, GA) at a dose rate of 5.519 Gy/min for the time required to apply the required dose. The X-ray beam was operated at 300kV and hardened using a 4mm Be, 3mm Al and 1.5mm Cu filter, and calibrated using NIST-traceable dosimetry. Radiation was delivered as a single dose of 4 or 8 Gy and controls (0 Gy) were sham irradiated.

### Sphere Formation Assays (SFAs) and Extreme Limiting Dilution Analysis (ELDA) Assays

For SFAs, cells were serially diluted in non-tissue culture treated 96-well plates under serum-free conditions. Starting cell densities for the serial dilution were as follows: 512 cells/well for 0 and 2 Gy, 1024 cells/well for 4 Gy, and 2048 cells/well for 8 Gy. After overnight incubation at 37°C, cells were treated and irradiated 1h later with 0, 2, 4, or 8 Gy using an X-ray irradiator. Cells were grown for 7-10 days and supplemented with 10 µl/well of GBM media with growth factors every other day. Upon formation of visibly distinct spheres, spheres formed in each well were counted using a conventional microscope and recorded. Four treatments were included in each plate resulting in an effective 20 wells/treatment condition. The number of spheres formed at each condition was adjusted for the number of cells plated per well and reported as a linear range average. This was subsequently normalized against the equivalent value for the non-irradiated control (DMSO/HyClone water, 0 Gy). The resulting data was plotted to determine the effect of the different treatment interventions on the sphere forming capacity (SFC) of the cells.

For ELDAs, glioma spheres were seeded at clonal densities under serum-free conditions in non-tissue culture treated 96-well plates using a serial dilution across the entire plate. Cells were treated, irradiated, supplemented, grown, counted, and SFC was determined as described for SFAs. For stem cell frequency calculations, open-access ELDA software [11] was used. Reported values were reported as a range (including lower, estimate and upper limits) as the reciprocal of stem cell frequency. Estimates were used for converting data into percent stem cell frequencies and results were plotted.

### Reprogramming Assay (GIC induction assay)

We used a fluorescence reporter system, with a vector coding for a ZsGreen (ZsG) fluorescent reporter protein fused to the carboxyl-terminal degron of ornithine decarboxylase (cODC). Briefly, cODC, an amino acid sequence recognized by the 26S proteasome leads to the immediate destruction of the associated fused protein. As a result, cells transfected with this vector and having normal 26S proteasome activity show no fluorescence, while cells with low proteasome activity accumulate the ZsG protein and fluoresce. Using HK-374 ZsG-cODC expressing cells grown as monolayers, single cell suspensions were sorted for the ZsG-negative population and collected using a BD FACSAria^TM^ III sorter (BD Biosciences, Franklin Lakes, NJ). Sorted cells were plated (∼27k cells/well) in 6-well tissue culture treated plates and incubated overnight. The following day, media was removed from the wells and cells were supplemented with fresh media containing the required drug treatments (duplicates or triplicates of each). One hour post-treatment plates were irradiated with 0 or 4 Gy and subsequently cultured for 5 days. On day 5 post-irradiation, cells were harvested and analyzed by flow cytometry (LSRFortessa^TM^ Cell Analyzer; BD Biosciences) for measuring green fluorescence through the Alexa Fluor 488 detector. Data collected was analyzed using FlowJo software (BD Biosciences). Non-ZsG transfected HK-374 cells were used as a background control for setting the required gates for our analysis. Fold increase in ZsG-positive cells were determined by normalizing the reported % of ZsG-positive cells in each sample to the average % ZsG-positive cells reported for the 0 Gy control treated wells.

### RT-PCR

Total RNA was isolated using TRIzol Reagent (Invitrogen, Waltham, MA, USA). cDNA synthesis was carried out using SuperScript Reverse Transcription IV (Invitrogen). Quantitative PCR was performed using a QuantStudio^TM^ Real-Time PCR System (Applied Biosystems, Carlsbad, CA, USA) using PowerUp^TM^SYBR^TM^ Green Master Mix (Applied Biosystems). C_t_ for each gene was determined after normalization to GAPDH or PPIA, and ΔΔC_t_ was calculated relative to the designated reference sample. Gene expression values were then reported as fold changes (2^-ΔΔCt^) as described by the manufacturer of the kit (Applied Biosystems). All PCR primers were synthesized by Invitrogen with GAPDH or PPIA being used as housekeeping genes. All primer sequences can be found in **Supplementary Table 2**.

### Bulk RNAseq

Raw data counts obtained from RNA sequencing of total RNA isolated from HK-374 monolayer cells 48h post-irradiation (treatment groups used: 0Gy_DMSO, 4Gy_DMSO and 4Gy_H151 1μM) by Novogene (Sacramento, CA), were input directly into iDEP [12], for performing differential gene expression and pathway analyses. Data was originally pre-processed using the following parameters: minimum counts per million (CPM) of 0.5, in 1 library and with a transformation of log2 (CPM+4) used for clustering and principal component analysis (PCA) to ensure sample quality. DESeq2 was used for identifying differentially expressed genes (DEGs) with the following parameters: false discovery rate (FDR) cutoff of 0.1 and minimum fold change of 2. Pathway enrichment analysis was done using the hallmarks.MSigDB geneset.

### Statistics

Apart from figures/tables obtained from iDEP, all other graphical representations and statistical analyses were performed using GraphPad Prism version 9. Unless otherwise specified, data was represented as mean +/- standard error mean (SEM) of at least 3 biological replicates. Statistical significance for one- and two-way ANOVA multiple comparisons testing was determined as follows: p<0.05 (*), p<0.01 (**), p<0.001 (***) and p <0.0001 (****)

## Results

### Identification of radiation-induced differentially expressed DAMPs and PRRs

To investigate the response of GBM cells to radiation, patient-derived HK-374 cells were sorted for non-stem glioma cells using our previously published imaging system for glioma-initiating cells (GICs), irradiated with 0 or 4 Gy and subjected to bulk RNAseq. At 48h post-IR we accessed differential gene expression of DAMPs and their respective PRRs. This time point is when there is maximal open chromatin in the promoter region of developmental TFs in response to radiation [8]. The unirradiated (0 Gy) and irradiated control (IR, 4 Gy) datasets of DEGs were searched against lists provided by Roh et al. [13] and the log2 fold changes of the identified DAMPs and PRRs were plotted as a heatmap. Radiation caused a specific expression pattern of DAMPs and PRRs that included up-regulation of TLR4 (**Figure 1**).

**Figure 1:**
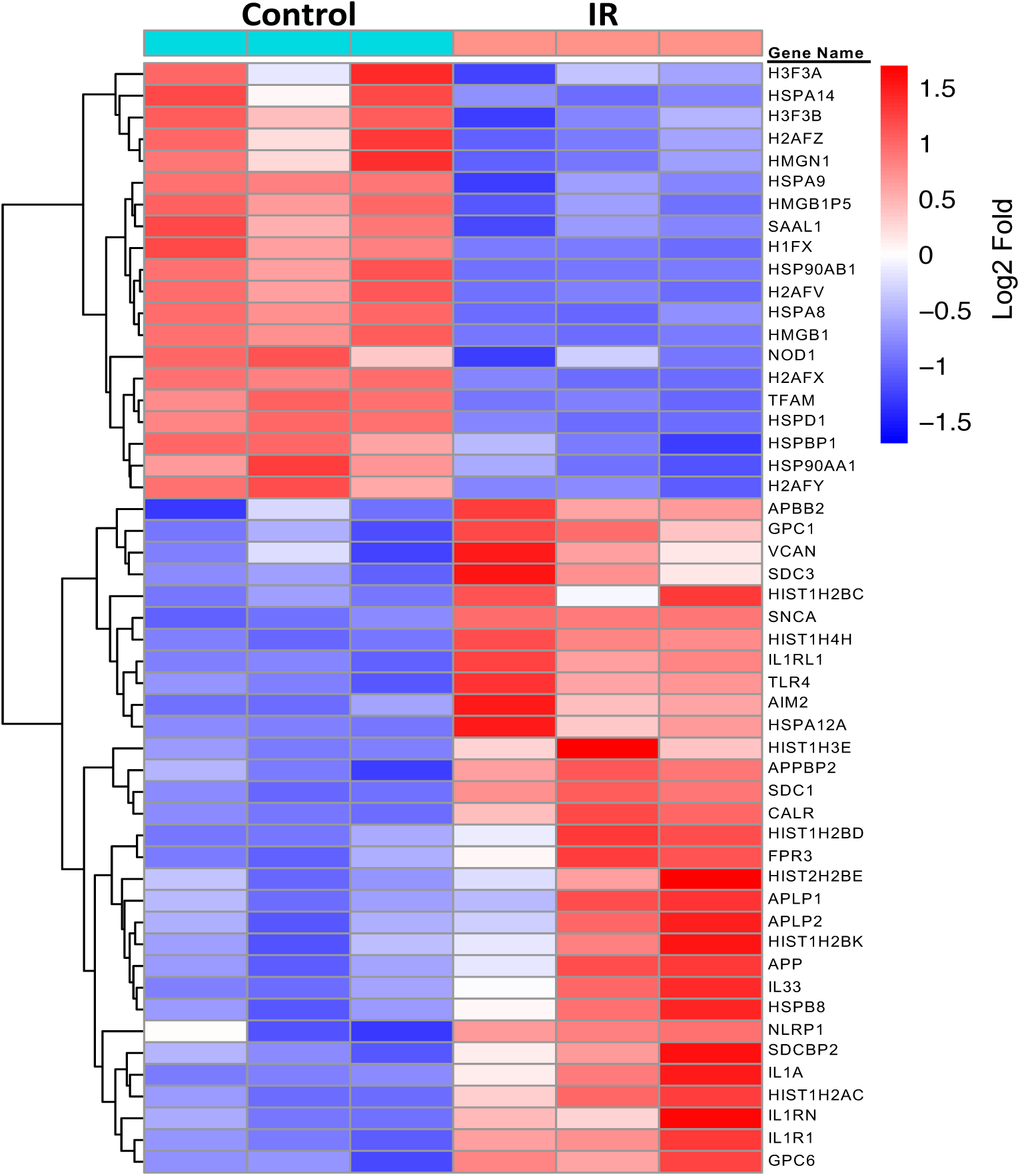
Radiation-induced expression of DAMPs and PRR genes in GBM. Heatmap representation of differentially expressed genes from an RNA sequencing analysis performed in HK-374 monolayer cells, 48h post 4 Gy irradiation. Data plotted as log2 fold changes. Control refers to unirradiated samples (0 Gy) and irradiation (IR) to 4 Gy.

Since TLR signaling has been implicated in cellular reprogramming, we initially focused on TLR4, upregulated by IR, and TLR3, which had been previously implicated in reprogramming [2]. Using a pharmacological approach, we evaluated the involvement of TLR3 and TLR4 in the self-renewal capacity of GBM cells using sphere formation assays (SFAs) and their effect on the number of GICs using extreme limiting dilution analysis (ELDA) assays.

TLRs generally signal through MyD88-dependent or -independent means. To test if radiation-induced reprogramming signals through a MyD88-dependent or -independent manner, we used specific inhibitors for TLR3 and TLR4 as well as chloroquine (CQ), a MyD88 inhibitor. TLR4 is capable of both types of signaling, while TLR3 exclusively mediates effects in a MyD88-independent manner [14]. However, there is evidence for CQ also inhibiting endocytic TLRs (TLR3/7/8/9) [15, 16], RIG-1, cGAS/STING, and processes such as autophagy [17].

In control samples, sphere formation capacity was decreased by more than 50% following irradiation with 4 Gy and almost 80% after 8 Gy (**Figure 2**). Treatment with TLR3i alone or in combination with radiation did not lead to a significant decrease in the SFC of the cells relative to the controls. Treatment with TLR4i alone led to a dose-dependent decrease in SFC compared to unirradiated control cells. This effect was also seen in combination with radiation (**Figure 2A**). A complete list of the numerical results is provided in **Supplementary Table 3**. Treatment with CQ alone led to a dose-dependent loss of SFC relative to the unirradiated control. When combined with 4 Gy, CQ showed a trend for a dose-dependent reduction in sphere formation, but these effects did not reach statistical significance (**Figure 2B**).

**Figure 2:**
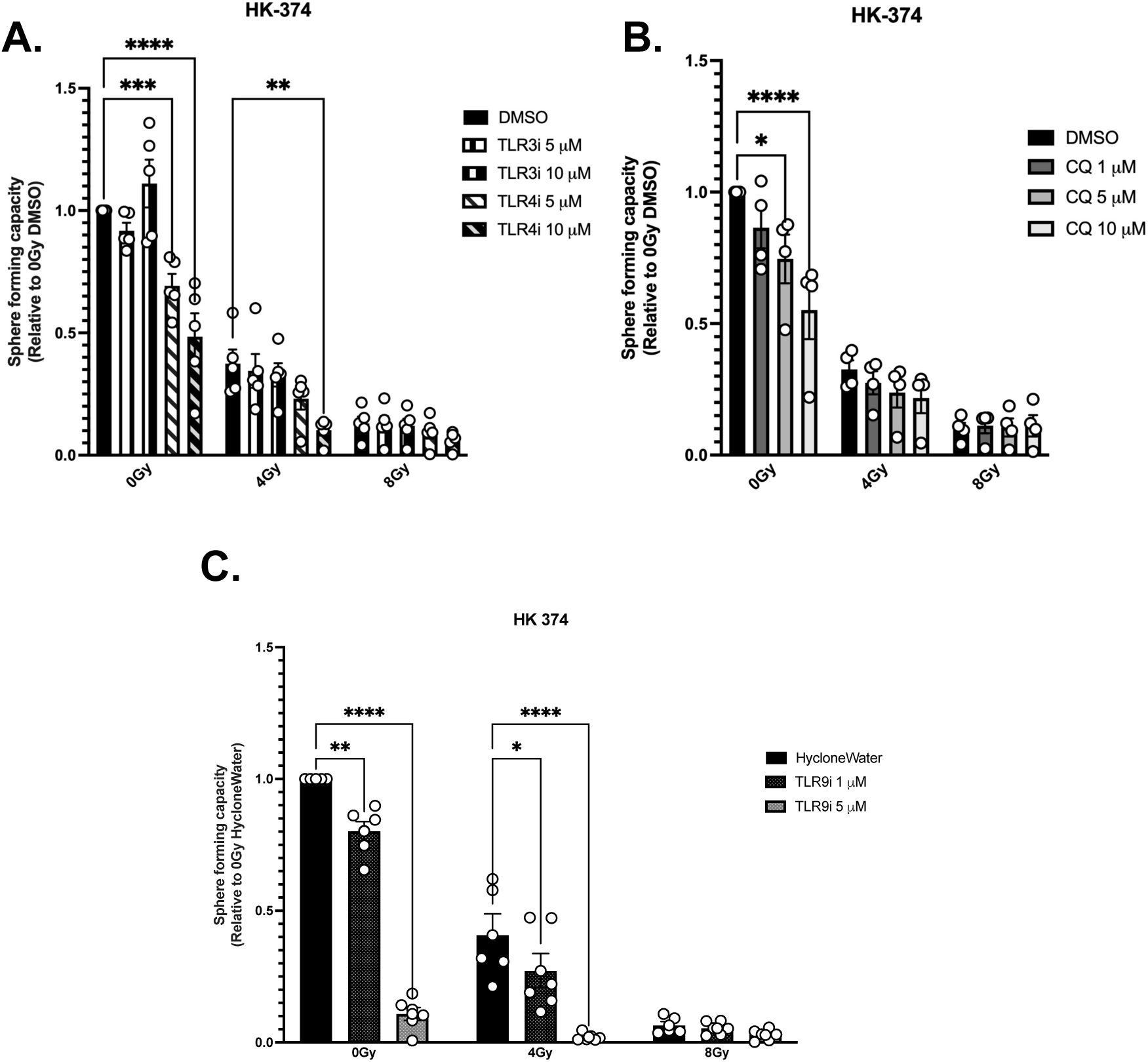
Combined radiation and TLR4/9, but not TLR3 inhibition reduces self-renewal capacity in GICs. In patient-derived HK-374 glioma spheres, **A.** sphere forming capacity (SFC) in presence of TLR3 and TLR4 inhibitors, n=5 biologically independent repeats. **B.** SFC in presence of chloroquine (CQ), n=4 biologically independent repeats. **C.** SFC in presence of a TLR9 inhibitor withn=5 for Hyclonewater (control), and n=6 for TLR9i at 1 and 5 μM. All data represented as mean +/- SEM. *p*-values were calculated using Two-way ANOVA. * *p*-value < 0.05, *\**\* *p*-value < 0.01, *** *p*-value < 0.001, **** *p*-value < 0.0001.

Our findings suggest that signaling through TLR4, but not TLR3, is involved in the maintenance of the self-renewal capacity of pre-existing GICs. Because CQ showed a trend for decreasing sphere formation while inhibition of TLR3 had no effect, we next tested if endocytic TLR9 [15] might be involved in mediating the effects of radiation on cellular plasticity, which had previously been reported to affect GIC maintenance and growth [18]. In SFAs using HK-374 cells 4 Gy in combination with TLR9i at the three highest concentrations tested, significantly decreased SFC relative to the 4 Gy control samples. (**Figure 2C**). A comprehensive list of the results is provided in **Supplementary Table 4**. At 8 Gy the number of countable spheres was in the single digits, which made detecting drug effects with statistical significance impossible. However, we included 8 Gy results for completeness. These findings highlight the critical role of TLR9 in GIC maintenance and suggest that TLR9 may also influence the self-renewal capacity of pre-existing GICs following radiation.

Since our data suggested TLR4 and TLR9 involvement in stem cell maintenance following irradiation, we used ELDA assays, a more rigorous approach, to evaluate effects of select receptor inhibitors on SFC and stem cell frequency. TLR9i was tested in cells from HK-374 (classical) and HK-308 (mesenchymal) glioma spheres (**Figure 3**). A significant, dose-dependent decrease in SFC was seen for both cell lines (**Figure 3A**). Detailed numerical results are provided in **Supplementary Table 5**. Consistent with the loss of self-renewal capacity, TLR9i treatment decreased the stem cell frequency in a dose-dependent manner (**Figure 3B**). This was statistically significant for HK-374 cells but not for HK-308 cells (**Supplementary Table 6**). Since both TLR4i and TLR9i affect GIC self-renewal, but only TLR9i affects the GIC frequency, this suggests that signaling through these receptors could be differentially affecting GICs following IR.

**Figure 3:**
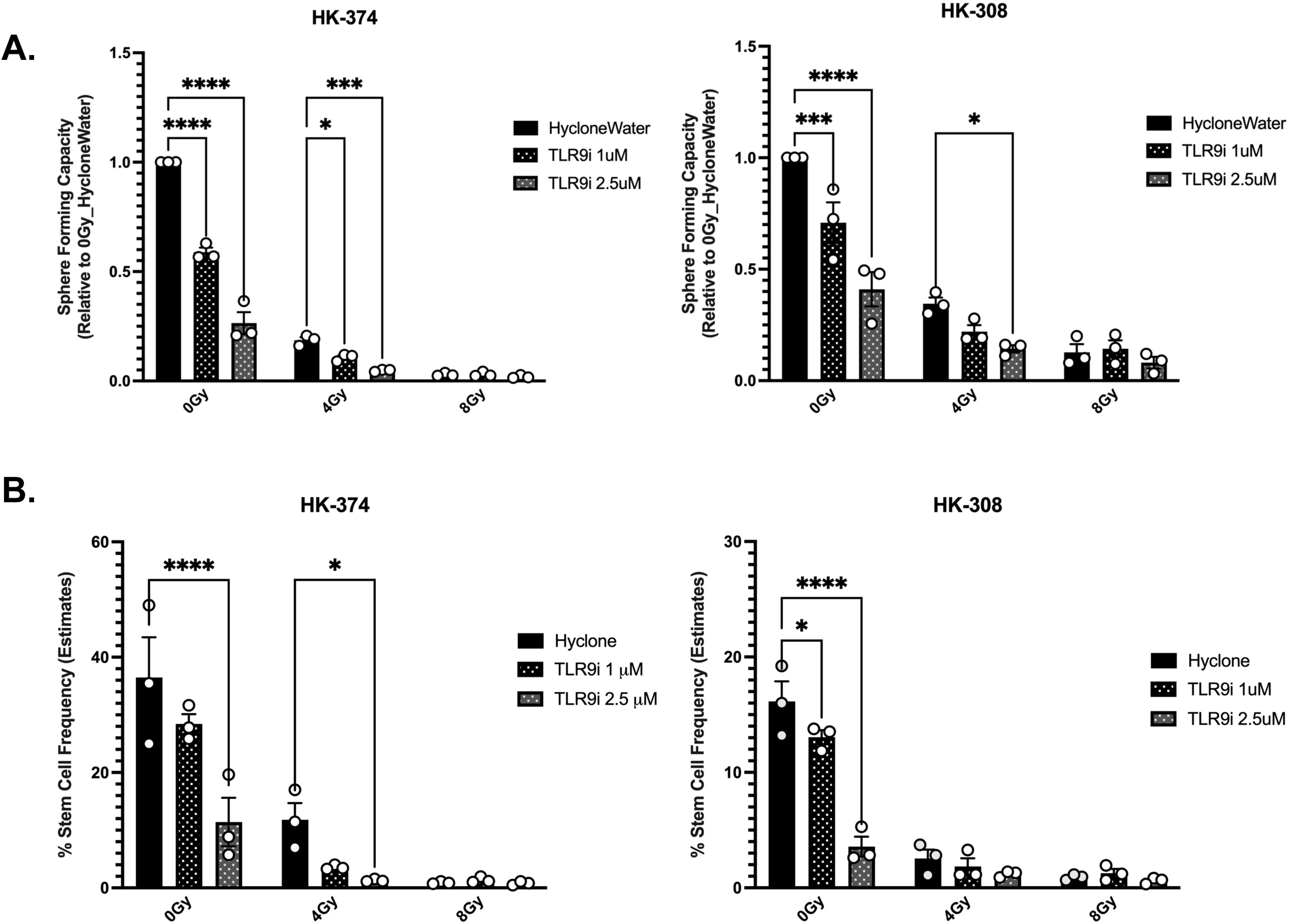
Combined radiation and TLR9 inhibition reduce self-renewal capacity and stem cell frequency in GICs. In patient-derived HK-374 and HK-308 glioma spheres **A.** SFC, n=3 biologically independent repeats. **B.** Stem cell frequency, n=3 biologically independent repeats. All data represented as mean +/- SEM. *p*-values were calculated using Two-way ANOVA. * *p*-value < 0.05, *** *p*-value < 0.001, **** *p*-value < 0.0001.

### TLR effects in radiation-induced induction of stemness

We next wanted to investigate the formation of new GICs from non-stem glioma cell populations following irradiation [7, 8] using reprogramming assays. We had previously shown that GICs have low 26S proteasome activity and developed a fluorescent reporter system to distinguish between the two populations (GICs and non-stem glioma cells) [19]. Sorted ZsG-negative non-stem glioma cells were irradiated. Five days later, the number of induced ZsG-positive GICs was assessed by FACS. Irradiation with a single dose of 4 Gy significantly increased the number of ZsG-positive cells relative to the sham irradiated control (**Figure 4A**), confirming our previous findings of *de novo* induction of GICs through irradiation [7, 8, 19]. Next, we tested if TLR3i, TLR4i or CQ would prevent radiation-induced plasticity after exposure to 4 Gy. Treatment with TLR3i (10 μM) led a significant decrease in the ZsG-positive population induced by 4 Gy. Likewise, CQ in combination with radiation, also reduced induction of ZsG-positive cells after 4 Gy at all concentrations (**Figure 4A**). Combination treatment of TLR4i with radiation was not significantly different from the irradiated control (**Supplementary Table 7**). These results suggest that radiation-induced phenotype conversion of non-GICs into GICs is likely dependent on TLR3, but not TLR4.

**Figure 4:**
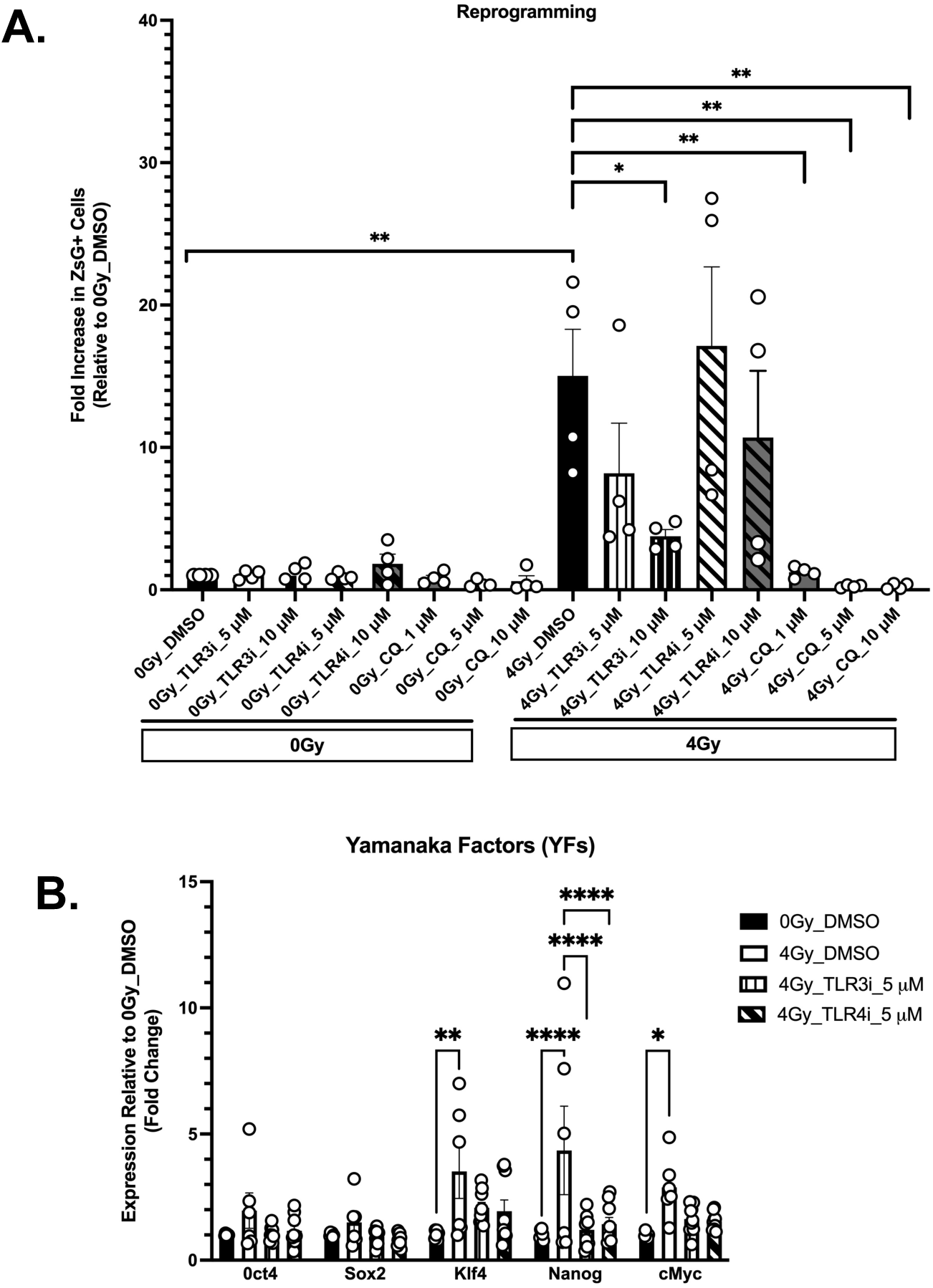
TLR3, but not TLR4 inhibition prevents radiation-induced phenotype conversion. **A.** Reprogramming assay in HK-374 monolayer cells treated with TLR3 and TLR4 inhibitors and CQ, n=4 biologically independent repeats for all conditions. **B.** Relative expression of Yamanaka factors in HK-374 monolayers (ZsG-negative sorted cells) irradiated with 4 Gy and treated with TLR3/TLR4 inhibitors, n=6 for 0/4 Gy, control and n=9 for 0/4 Gy, 5 μM TLR3/4i biologically independent repeats for each condition. All data represented as mean +/- SEM. *p*-values were calculated using One-way ANOVA for **A**, Two-way ANOVA for **B**. * *p*-value < 0.05, *\**\* *p*-value < 0.01, **** *p*-value < 0.0001.

We next tested if treatment of sorted non-stem glioma cells with TLR3 and TLR4 inhibitors in combination with radiation would lead to changes in the expression of YFs. For our experiments, changes in YF and Nanog gene expression levels in sorted non-stem glioma cells were assessed by RT-PCR, 48h after IR with 4 Gy and treatment with TLR3i or TLR4i. We observed increases in the expression of all YFs and Nanog that reached statistical significance for Klf4, Nanog and c-Myc (**Figure 4B**), confirming data previously reported [7, 8]. The addition of TLR3i or TLR4i prevented or attenuated the effect of radiation on the expression of YFs and Nanog (**Figure 4B**). Complete two-way ANOVA analysis results can be found in **Supplementary Table 8**. These findings suggested that TLR3 and/or TLR4 might be involved in the process of radiation-induced phenotype conversion.

### Role of cGAS/STING signaling pathway in radiation-induced cellular plasticity

Radiation exposure is known to lead to DNA breaks. The resulting fragments of free dsDNA can be considered DAMPs. As such, the potential role of the cGAS/STING pathway, a signaling pathway involved in the sensing of free cytosolic DNA [20], was evaluated. First, we tested if our glioma lines expressed cGAS/STING using one set of human specific primers for cGAS (hcGAS) and two different primer sets for STING expression (hSTING_1/2) (**Figure 5A**). To determine whether the cGAS/STING pathway could be playing a role in radiation-induced cellular plasticity we performed ELDA assays using select human cGAS (G140) and STING (H151) inhibitors and evaluated their effects on SFC and stem cell frequency (**Figure 5B-D**). In HK-374 cells, cGAS inhibition did not decrease sphere formation or stem cell frequencies (**Figure 5B, Supplementary Table 9**). In contrast, STINGi combined with radiation resulted in a statistically significant dose-dependent decrease in SFC of HK-374 and HK-157 cells. HK-308 cells only showed a significant, dose-dependent decrease in SFC in non-irradiated samples (**Figure 5C**). This suggested that inhibition of steady state STING signaling is affecting stem cell self-renewal, proliferation and/or survival. Complete overview of two-way ANOVA multiple comparisons analysis can be found in **Supplementary Tables 10-11**. Analysis of the changes in stem cell frequencies of ELDA assays treated with the STINGi showed a significant decrease in stem cell frequency at the highest STINGi concentration in combination with 2 Gy irradiation for HK-374 cells, while such a decrease was only evident in the absence of radiation for the other two cell lines tested (**Figure 5D**, **Supplementary Tables 12-15**). These results indicate a role for non-canonical STING signaling in radiation-induced cellular plasticity.

**Figure 5:**
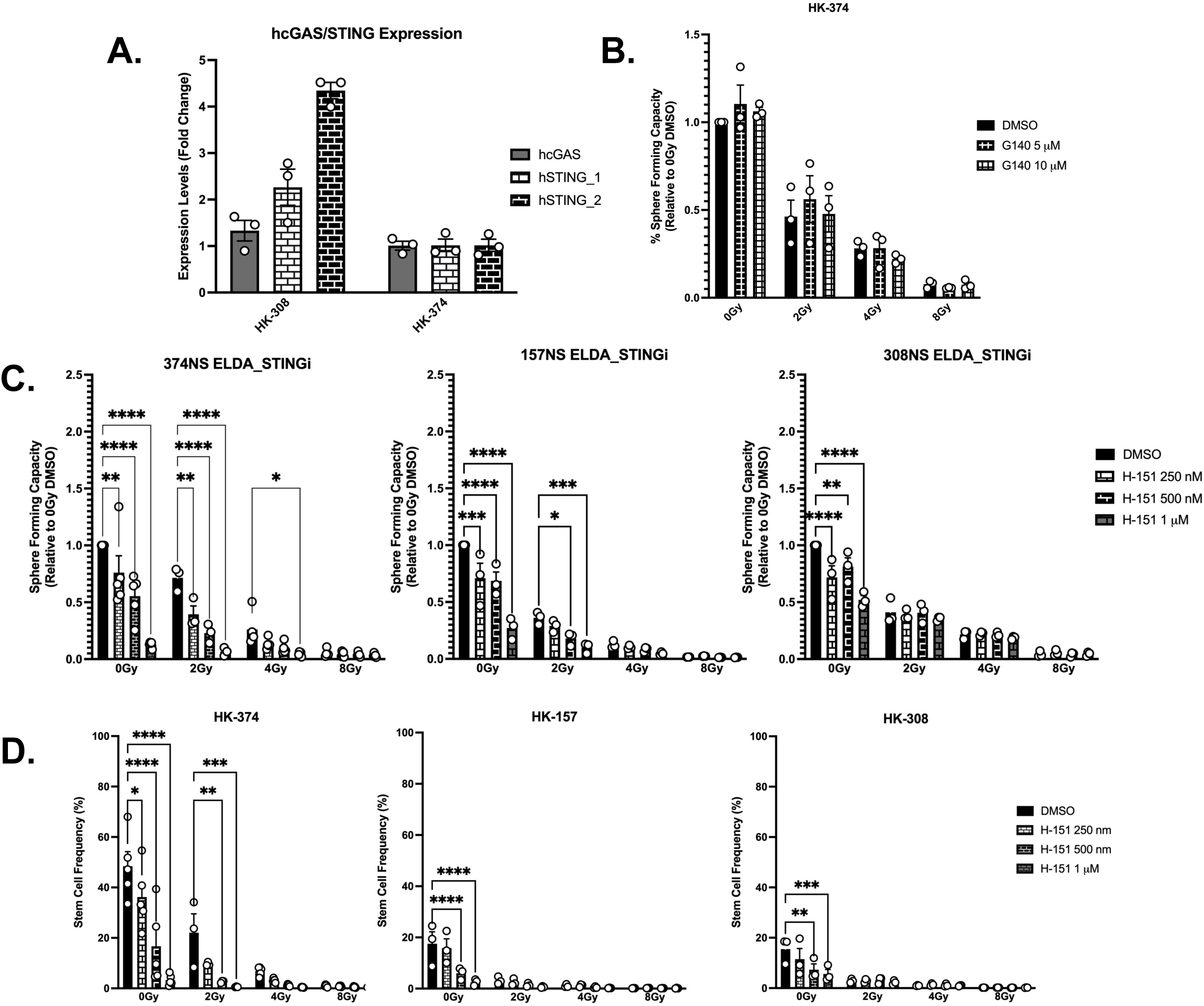
STING inhibition differentially affect self-renewal capacity and stem cell frequency in GICs. **A.** cGAS and STING expression in patient-derived HK-374 and HK-308. **B.** SFC in patient-derived HK-374, HK-157, HK-308 glioma treated with hcGASi (G140), n=3 biologically independent repeats. **C.** SFC for HK-374, HK-157, and HK-308 glioma spheres treated with human STINGi (H-151). For HK-374, n=5 biologically independent repeats for 0/4/8 Gy and n=3 for 2 Gy. For HK-157 and HK-308, n=3 biologically independent repeats for all conditions. **D.** Stem cell frequency in HK-374, HK-157, and HK-308 glioma spheres treated with H-151. For HK-374 n=5 biologically independent repeats for 0/4/8 Gy, n=3 for 2 Gy. For HK-157 and HK-308, n=3 independent biologically repeats for all conditions tested. All data represented as mean +/- SEM. *p*-values were calculated using Two-way ANOVA. * *p*-value < 0.05, *\**\* *p*-value < 0.01, *** *p*-value < 0.001, **** *p*-value < 0.0001.

To further analyze the effects of STING inhibition at the transcriptomic level, bulk RNAseq analysis was performed in HK-374 cells, 48h after irradiation with 4 Gy. Differential gene expression analysis identified 145 genes and 1853 genes upregulated by 4 Gy + STINGi (denoted as IR + STINGi) relative to IR alone and unirradiated (DMSO, 0 Gy) samples. 1937 genes and 2688 genes were downregulated in the combination treatment group relative to IR and unirradiated control, respectively. Pathway analysis for DEGs identified from RNAseq (**Figure 6A**) for IR *vs* control conditions showed downregulation of the hallmark gene sets for *G2M checkpoint* and *E2F targets,* which was in accordance with expected radiation effects on cell cycle and DNA replication pathways. Among gene sets shown to be enriched in the IR group, *epithelial mesenchymal transition*, *KRAS signaling*, *p53 pathway*, *coagulation*, *inflammatory response*, *apoptosis*, *TNFα signaling via NFκB*, *angiogenesis*, *complement* and *hypoxia* were identified. Upregulation of these gene sets aligned with the known cytotoxic and pro-inflammatory effects of radiation.

**Figure 6:**
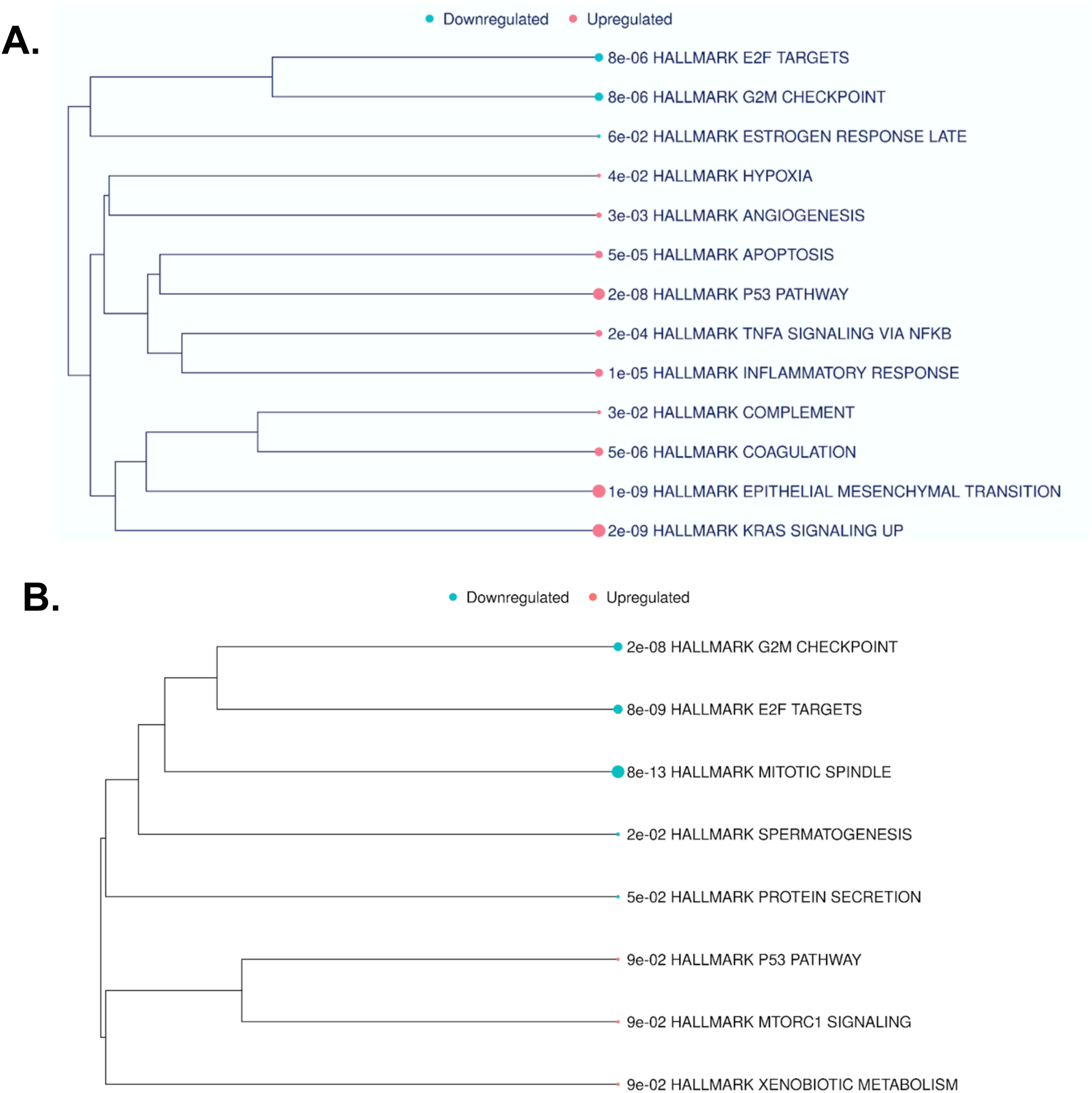
Pathway Analysis of radiation-induced DEGs in GBM. RNA sequencing data pathway analysis of **A**. 4 Gy versus control (0 Gy) and **B.** 4 Gy + STINGi (1 μM H151) versus 4 Gy in HK-374 monolayer samples 48h post-irradiation.

Performing a similar pathway analysis for the DEGs from the IR + STINGi *vs* IR (**Figure 6B)** identified many of the same downregulated gene sets as those listed above, with the addition of *mitotic spindle*, *spermatogenesis* and *protein secretion*. *mTORC1 signaling*, *xenobiotic metabolism* and *p53 pathway* were found to be upregulated in the IR + STINGi group relative to 4 Gy.

## Discussion

In this study, we show that GIC maintenance after irradiation is mediated non-canonical STING, TLR4, and TLR9 signaling, while radiation-induced *de novo* GIC induction involves TLR3. These insights reveal a key role for PRR signaling in radiation-induced cellular plasticity, hinting at potential mechanisms of GBM therapy resistance and progression.

Our GBM RNAseq dataset (**Figure 1A**) revealed several differentially expressed DAMPs and PRRs following irradiation. However, HMGB1, a common DAMP released post-irradiation [21], was shown to be slightly downregulated compared to unirradiated controls. This discrepancy could be due to our experimental conditions such as time point (48h), radiation dose, and/or the cell line used. Since RNAseq is limited to gene expression changes, nucleic acid DAMPs such as those recognized by cGAS/STING, endocytic TLRs, and RLRs [22] might have been missed. Additionally, receptors that do not show significant mRNA level changes, and thus not identified in the list of DEGs, could be contributing to radiation-induced cellular plasticity [23].

In sphere-forming capacity assays we show that stem cell maintenance and self-renewal following radiation likely involve TLR4 and TLR9 signaling (**Figures 2-3**). The TLR3 inhibitor showed no effect on SFC, implicating a MyD88-dependent pathway in RT-induced GIC maintenance. Since both TLR4 and TLR9 inhibition affect GIC self-renewal, yet only one reduces GIC frequency (**Figure 3**), these receptors may mediate radiation-induced cellular plasticity effects through different mechanisms. In contrast with the literature showing GICs downregulating TLR4 as a means of maintaining their self-renewal capabilities [24], our results showed a decrease in SFC following TLR4i treatment. In one study, signaling through the TLR4-TBK1 axis (MyD88-independent) leads to the inhibition of retinoblastoma binding protein 5 (RBBP5), a TF elevated in GICs, and shown to be necessary for their maintenance [24]. Downregulation of TLR4 in in this context removes the inhibitory effect of TBK1 on RBBP5, allowing the expression of pluripotency genes and subsequent stem cell maintenance. The discrepancy with our present findings could be explained by cell line heterogeneity of cell lines used and/or broader effects of the TLR4 inhibitor. Nevertheless, both studies highlight a role of the MyD88-dependent pathway in GIC maintenance and self-renewal.

In our CQ experiments (**Figure 2B**), we observed a dose-dependent reduction in SFC, both without radiation and at 4 Gy (**Figures 2C**). The broad effects of CQ could explain the differences between TLR3/9 and CQ findings. CQ inhibits endosomal TLRs including TLR3 and TLR9 as well as the cGAS/STING, and RIG-1/MAVS signaling pathways. CQ further affects MyD88 signaling [16] and impairs autophagic machinery [25] which is implicated in reprogramming [26]. We observed a trend for CQ decreasing GIC self-renewal capacity and inhibited reprogramming more effectively than TLR3 inhibition. We also did not observe a statistically significant decrease following irradiation like in our TLR9i findings. This supports the potential role of TLR9 in radiation-induced cellular plasticity and suggests that reprogramming relies on multiple CQ-targeted pathways [2, 17, 25, 27-30]. While a single receptor may mediate radiation-induced reprogramming, our CQ data indicate that multiple receptors, pathways, and/or processes are likely involved. Further, RT effects evaluated by SFAs/ELDAs and reprogramming assays do not necessarily diverge on the same receptors (TLR4/TLR9 versus TLR3). The respective assays evaluate two distinct processes; SFAs/ELDAs evaluate the formation of new GICs from existing GIC enriched populations, while reprogramming assays look at newly generated GICs from a previously GIC-exhausted population. Nevertheless, both processes fall under the umbrella of cellular plasticity, hinting at the complexity of the radiation-induced cellular plasticity response.

Several distinct molecules functioning as DAMPs can be released following IR, including cytosolic dsDNA fragments, micronuclei, and nucleotides [21], which are key signals for cellular stress. Each molecule can engage either a single, unique receptor or a combination of different receptors [22], and in doing so activate important downstream signaling cascades. Our ELDA data suggest radiation-induced cellular plasticity occurs through cGAS-independent but STING-dependent signaling (**Figure 5**). This aligns with evidence for non-canonical STING activation, independent of cGAS [31-33] and highlight a novel possibility for cellular plasticity events following irradiation. Given the magnitude of effects seen following STING inhibition in our ELDAs, future reprogramming assays using cGAS/STING inhibitors are warranted.

Inhibition effects for TLR4 and TLR9 were consistent across classical (HK-374) and mesenchymal (HK-308/345) lines, while STINGi showed stronger effects in classical and proneural (HK-157) compared to mesenchymal (HK-308) lines. Additional experiments in each subtype in future studies will determine if there is indeed a preferential signaling pathway being engaged following irradiation. Regardless, TLR4, TLR9 and STING inhibitors reduced SFC suggesting receptor crosstalk mediates radiation responses rather than a single unique receptor and/or pathway. Recognition of the same DAMP by multiple receptors enables synergistic downstream action characterized by the activation of multiple signaling pathways. Further investigation into downstream signaling is required to understand how these processes are coordinated during radiation-induced cellular plasticity events.

## Conclusion

Taken together, this study identifies critical pathways and receptors involved in radiation-induced cellular plasticity in GBM, emphasizing the roles of cGAS-independent STING signaling, TLR4, TLR9, and autophagy-related processes. *De novo* stemness induction implicates TLR3, with CQ experiments suggesting additional pathways and receptor crosstalk. Our findings highlight the interplay between DAMPs, PRRs, and downstream signaling. These insights advance understanding of radiation-induced resistance and progression mechanisms, offering a foundation for targeting plasticity-related pathways in future GBM therapeutic strategies.

## Data Availability Statement

All data are available from the corresponding author upon reasonable request.

## Author contributions

FP consived of the study. AI performed the experiments, AMC, LA, IA, and FP drafted the paper.

All authors wrote the final manuscript

