## Supplemental Materials for "Targeting Innate Immune Signaling in Glioma-Initiating Cells Impairs Self-Renewal and Radiation-Induced Cellular Plasticity"

**Supplementary Tables and Figures:**

**Supplementary Table 1.** List of inhibitors used and their specific characteristics.

| Target | Inhibitor name | Abbreviations within text | Manufacturer | Cat # | Function |
|---|---|---|---|---|---|
| TLR3 | TLR3/dsRNA complex inhibitor | TLR3i | Calbiochem® | 614310 | Directly and competitively inhibit dsRNA binding to TLR3 |
| TLR4 | TAK 242 | TLR4i | Calbiochem® | 614316 | Disrupts TLR4 interaction with downstream adaptor molecules TIRAP and TRAM (affecting both NFκB and IFN signaling) |
| TLR9 | E6446 dihydrochloride | TLR9i | Selleck Chemicals | S6719 | Specific TLR9 inhibitor shown to disrupt CpG oligonucleotide induced NFκB signaling |
| cGAS | Human cGAS inhibitor | cGASi | InvivoGen | inh-g140 | Targets the cGAS catalytic pocket, preventing the synthesis of downstream messenger 2', 3'-cGAMP |
| STING | H-151 | STINGi, H151 | MCE | HY-112693 | Decreases TBK1 phosphorylation and suppresses STING palmitoylation (required for STING activation of type I interferon response) |
| MyD88 & endosomal TLRs | Chloroquine diphosphate salt | CQ | Sigma-Aldrich | C6628 | Inhibition of endosomal TLRs and MyD88 signaling by decreasing IRAK4 and IRF7 molecules and inhibiting IFNa synthesis; cGAS/STING, RIG-1 signaling and autophagy have also been shown to be affected by CQ [63] |

**Supplementary Table 2.** Primer sequences (Integrated DNA Technologies and Origene) used for RT-PCR experiments Note: designation “h” indicate human specific primer.

| | |
|---|---|
| Oct4 | |
| Forward | CATAGTCGCTGCTTGATCGCTTG |
| Reverse | GAGAACCGAGTGAGAGGCAACC |
| Sox2 | |
| Forward | TTGCGTGAGTGTGGATGGGATTGGTG |
| Reverse | GGGAAATGGGAGGGGTGCAAAAGAGG |
| Klf4 | |
| Forward | GGTCCGACCTGGAAAATGCT |
| Reverse | ACCAGGCACTACCGTAAACACA |
| Nanog | |
| Forward | TGCGTCACACCATTGCTATTCTTC |
| Reverse | AATACCTCAGCCTCCAGCAGATG |
| cMyc | |
| Forward | CACTGTCCAACTTGACCCTCTTG |
| Reverse | CGTCTCCACACATCAGCACAA |
| hcGAS | Integrated DNA Technologies |
| Forward | AGGAAGCAACTACGACTAAAGC |
| Reverse | TCACAGCACGTTTTAGATTTTCC |
| hSTING_1 | Integrated DNA Technologies |
| Forward | TCAAGGATCGGGTTTACAGC |
| Reverse | GCTTGACTGTATTGTGACATGG |
| hSTING_2 | |
| Forward | CCTGAGTCTCAGAACAACTGCC |
| Reverse | GGTCTTCAAGCTGCCCACAGTA |

**Supplementary Table 3.** Two-way ANOVA analysis results of multiple comparisons for data represented in Figure 2A (SFAs of TLR3/4i).

| Dunnett's multiple comparisons test | Mean Diff. | 95.00% CI of diff. | Summary | Adjusted P Value |
|---|---|---|---|---|
| **0Gy** | | | | |
| DMSO vs. TLR3i 5uM | 0.08196 | -0.09831 to 0.2622 | ns | 0.6137 |
| DMSO vs. TLR3i 10uM | -0.1105 | -0.2908 to 0.06976 | ns | 0.3548 |
| DMSO vs. TLR4i 5uM | 0.3074 | 0.1272 to 0.4877 | *** | 0.0003 |
| DMSO vs. TLR4i 10uM | 0.5153 | 0.3350 to 0.6956 | **** | <0.0001 |
| **4Gy** | | | | |
| DMSO vs. TLR3i 5uM | 0.02929 | -0.1510 to 0.2096 | ns | 0.9828 |
| DMSO vs. TLR3i 10uM | 0.04546 | -0.1348 to 0.2257 | ns | 0.9214 |
| DMSO vs. TLR4i 5uM | 0.1427 | -0.03756 to 0.3230 | ns | 0.1585 |
| DMSO vs. TLR4i 10uM | 0.2720 | 0.09175 to 0.4523 | ** | 0.0014 |
| **8Gy** | | | | |
| DMSO vs. TLR3i 5uM | 0.003462 | -0.1768 to 0.1837 | ns | >0.9999 |
| DMSO vs. TLR3i 10uM | 0.01012 | -0.1701 to 0.1904 | ns | 0.9998 |
| DMSO vs. TLR4i 5uM | 0.03539 | -0.1449 to 0.2157 | ns | 0.9662 |
| DMSO vs. TLR4i 10uM | 0.06917 | -0.1111 to 0.2494 | ns | 0.7376 |

**Supplementary Table 4.** Two-way ANOVA analysis results of multiple comparisons for data represented in Figure 2C (SFA of TLR9i).

| Dunnett's multiple comparisons test | Predicted (LS) mean diff. | 95.00% CI of diff. | Summary | Adjusted P Value |
|---|---|---|---|---|
| **0Gy** | | | | |
| HycloneWater vs. TLR9i 1uM | 0.1983 | 0.05135 to 0.3452 | ** | 0.0041 |
| HycloneWater vs. TLR9i 2.5uM | 0.3365 | 0.1335 to 0.5395 | *** | 0.0003 |
| HycloneWater vs. TLR9i 5uM | 0.8921 | 0.7452 to 1.039 | **** | <0.0001 |
| HycloneWater vs. TLR9i 7.5uM | 1.000 | 0.7970 to 1.203 | **** | <0.0001 |
| HycloneWater vs. TLR9i 10uM | 1.000 | 0.7970 to 1.203 | **** | <0.0001 |
| **4Gy** | | | | |
| HycloneWater vs. TLR9i 1uM | 0.1354 | -0.01150 to 0.2823 | ns | 0.0814 |
| HycloneWater vs. TLR9i 2.5uM | 0.1711 | -0.03193 to 0.3741 | ns | 0.1287 |
| HycloneWater vs. TLR9i 5uM | 0.3871 | 0.2402 to 0.5341 | **** | <0.0001 |
| HycloneWater vs. TLR9i 7.5uM | 0.4061 | 0.2031 to 0.6091 | **** | <0.0001 |
| HycloneWater vs. TLR9i 10uM | 0.4070 | 0.2039 to 0.6100 | **** | <0.0001 |
| **8Gy** | | | | |
| HycloneWater vs. TLR9i 1uM | 0.01042 | -0.1365 to 0.1573 | ns | 0.9997 |
| HycloneWater vs. TLR9i 2.5uM | -0.005347 | -0.2083 to 0.1977 | ns | >0.9999 |
| HycloneWater vs. TLR9i 5uM | 0.03626 | -0.1107 to 0.1832 | ns | 0.9595 |
| HycloneWater vs. TLR9i 7.5uM | 0.06098 | -0.1420 to 0.2640 | ns | 0.9132 |
| HycloneWater vs. TLR9i 10uM | 0.06089 | -0.1421 to 0.2639 | ns | 0.9137 |

**Supplementary Table 5.** Two-way ANOVA analysis results of multiple comparisons for HK-374 & HK-308 gliomasphere data represented in Figure 3A (ELDA of TLR9i).

HK-374

| Dunnett's multiple comparisons test | Mean Diff. | 95.00% CI of diff. | Summary | Adjusted P Value |
|---|---|---|---|---|
| **0Gy** | | | | |
| HycloneWater vs. TLR9i 1uM | 0.4114 | 0.3463 to 0.4765 | **** | <0.0001 |
| HycloneWater vs. TLR9i 2.5uM | 0.7356 | 0.6705 to 0.8008 | **** | <0.0001 |
| **4Gy** | | | | |
| HycloneWater vs. TLR9i 1uM | 0.07849 | 0.01337 to 0.1436 | * | 0.0181 |
| HycloneWater vs. TLR9i 2.5uM | 0.1391 | 0.07395 to 0.2042 | *** | 0.0001 |
| **8Gy** | | | | |
| HycloneWater vs. TLR9i 1uM | -0.001270 | -0.06639 to 0.06385 | ns | 0.9984 |
| HycloneWater vs. TLR9i 2.5uM | 0.009709 | -0.05541 to 0.07483 | ns | 0.9116 |

HK-308

| Dunnett's multiple comparisons test | Mean Diff. | 95.00% CI of diff. | Summary | Adjusted P Value |
|---|---|---|---|---|
| **0Gy** | | | | |
| HycloneWater vs. TLR9i 1uM | 0.2913 | 0.1328 to 0.4498 | *** | 0.0007 |
| HycloneWater vs. TLR9i 2.5uM | 0.5899 | 0.4314 to 0.7485 | **** | <0.0001 |
| **4Gy** | | | | |
| HycloneWater vs. TLR9i 1uM | 0.1255 | -0.03302 to 0.2840 | ns | 0.1295 |
| HycloneWater vs. TLR9i 2.5uM | 0.2021 | 0.04364 to 0.3607 | * | 0.0127 |
| **8Gy** | | | | |
| HycloneWater vs. TLR9i 1uM | -0.01601 | -0.1745 to 0.1425 | ns | 0.9581 |
| HycloneWater vs. TLR9i 2.5uM | 0.04615 | -0.1124 to 0.2047 | ns | 0.7122 |

**Supplementary Table 6.** Two-way ANOVA analysis results of multiple comparisons for HK-374 & HK-308 gliomasphere data represented in Figure 3B (SC freq for TLR9i ELDA).

HK-374

| Dunnett's multiple comparisons test | Mean Diff. | 95.00% CI of diff. | Summary | Adjusted P Value |
|---|---|---|---|---|
| **0Gy** | | | | |
| Hyclone vs. TLR9i 1uM | 8.072 | -1.892 to 18.04 | ns | 0.1195 |
| Hyclone vs. TLR9i 2.5uM | 25.09 | 15.12 to 35.05 | **** | <0.0001 |
| **4Gy** | | | | |
| Hyclone vs. TLR9i 1uM | 8.243 | -1.721 to 18.21 | ns | 0.1109 |
| Hyclone vs. TLR9i 2.5uM | 10.53 | 0.5648 to 20.49 | * | 0.0380 |
| **8Gy** | | | | |
| Hyclone vs. TLR9i 1uM | -0.4670 | -10.43 to 9.498 | ns | 0.9908 |
| Hyclone vs. TLR9i 2.5uM | 0.09407 | -9.870 to 10.06 | ns | 0.9996 |

**HK_308**

| Dunnett's multiple comparisons test | Mean Diff. | 95.00% CI of diff. | Summary | Adjusted P Value |
|---|---|---|---|---|
| **0Gy** | | | | |
| Hyclone vs. TLR9i 1uM | 3.091 | 0.4562 to 5.725 | * | 0.0213 |
| Hyclone vs. TLR9i 2.5uM | 12.57 | 9.938 to 15.21 | **** | <0.0001 |
| **4Gy** | | | | |
| Hyclone vs. TLR9i 1uM | 0.6962 | -1.938 to 3.331 | ns | 0.7543 |
| Hyclone vs. TLR9i 2.5uM | 1.347 | -1.288 to 3.981 | ns | 0.3831 |
| **8Gy** | | | | |
| Hyclone vs. TLR9i 1uM | -0.3444 | -2.979 to 2.290 | ns | 0.9311 |
| Hyclone vs. TLR9i 2.5uM | 0.2987 | -2.336 to 2.933 | ns | 0.9476 |

**Supplementary Table 7.** One-way ANOVA analysis results of multiple comparisons for data represented in Figure 4A (Reprogramming assay using TLR3/4i and CQ).

| Tukey's multiple comparisons test | Mean Diff. | 95.00% CI of diff. | Summary | Adjusted P Value |
|---|---|---|---|---|
| 0Gy_DMSO vs. 0Gy_TLR3i_5uM | -0.02922 | -11.23 to 11.17 | ns | >0.9999 |
| 0Gy_DMSO vs. 0Gy_TLR3i_10uM | -0.2207 | -11.42 to 10.98 | ns | >0.9999 |
| 0Gy_DMSO vs. 0Gy_TLR4i_5uM | 0.08227 | -11.11 to 11.28 | ns | >0.9999 |
| 0Gy_DMSO vs. 0Gy_TLR4i_10uM | -0.8242 | -12.02 to 10.37 | ns | >0.9999 |
| 0Gy_DMSO vs. 0Gy_CQ_1uM | 0.1973 | -11.00 to 11.39 | ns | >0.9999 |
| 0Gy_DMSO vs. 0Gy_CQ_5uM | 0.5770 | -10.62 to 11.77 | ns | >0.9999 |
| 0Gy_DMSO vs. 0Gy_CQ_10uM | 0.3895 | -10.81 to 11.59 | ns | >0.9999 |
| 0Gy_DMSO vs. 4Gy_DMSO | -14.02 | -25.22 to -2.829 | ** | 0.0036 |
| 0Gy_DMSO vs. 4Gy_TLR3i_5uM | -7.188 | -18.38 to 4.008 | ns | 0.6122 |
| 0Gy_DMSO vs. 4Gy_TLR3i_10uM | -2.766 | -13.96 to 8.430 | ns | >0.9999 |
| 0Gy_DMSO vs. 4Gy_TLR4i_5uM | -16.13 | -27.33 to -4.936 | *** | 0.0004 |
| 0Gy_DMSO vs. 4Gy_TLR4i_10uM | -9.703 | -20.90 to 1.493 | ns | 0.1578 |
| 0Gy_DMSO vs. 4Gy_CQ_1uM | -0.2364 | -11.43 to 10.96 | ns | >0.9999 |
| 0Gy_DMSO vs. 4Gy_CQ_5uM | 0.7525 | -10.44 to 11.95 | ns | >0.9999 |
| 0Gy_DMSO vs. 4Gy_CQ_10uM | 0.7521 | -10.44 to 11.95 | ns | >0.9999 |
| 0Gy_TLR3i_5uM vs. 0Gy_TLR3i_10uM | -0.1914 | -11.39 to 11.00 | ns | >0.9999 |
| 0Gy_TLR3i_5uM vs. 0Gy_TLR4i_5uM | 0.1115 | -11.08 to 11.31 | ns | >0.9999 |
| 0Gy_TLR3i_5uM vs. 0Gy_TLR4i_10uM | -0.7950 | -11.99 to 10.40 | ns | >0.9999 |
| 0Gy_TLR3i_5uM vs. 0Gy_CQ_1uM | 0.2265 | -10.97 to 11.42 | ns | >0.9999 |
| 0Gy_TLR3i_5uM vs. 0Gy_CQ_5uM | 0.6063 | -10.59 to 11.80 | ns | >0.9999 |
| 0Gy_TLR3i_5uM vs. 0Gy_CQ_10uM | 0.4187 | -10.78 to 11.61 | ns | >0.9999 |
| 0Gy_TLR3i_5uM vs. 4Gy_DMSO | -14.00 | -25.19 to -2.800 | ** | 0.0037 |
| 0Gy_TLR3i_5uM vs. 4Gy_TLR3i_5uM | -7.159 | -18.35 to 4.037 | ns | 0.6186 |
| 0Gy_TLR3i_5uM vs. 4Gy_TLR3i_10uM | -2.737 | -13.93 to 8.459 | ns | >0.9999 |

| | | | | |
|---|---|---|---|---|
| 0Gy_TLR3i_5uM vs. 4Gy_TLR4i_5uM | -16.10 | -27.30 to -4.907 | *** | 0.0004 |
| 0Gy_TLR3i_5uM vs. 4Gy_TLR4i_10uM | -9.674 | -20.87 to 1.522 | ns | 0.1611 |
| 0Gy_TLR3i_5uM vs. 4Gy_CQ_1uM | -0.2071 | -11.40 to 10.99 | ns | >0.9999 |
| 0Gy_TLR3i_5uM vs. 4Gy_CQ_5uM | 0.7818 | -10.41 to 11.98 | ns | >0.9999 |
| 0Gy_TLR3i_5uM vs. 4Gy_CQ_10uM | 0.7813 | -10.41 to 11.98 | ns | >0.9999 |
| 0Gy_TLR3i_10uM vs. 0Gy_TLR4i_5uM | 0.3029 | -10.89 to 11.50 | ns | >0.9999 |
| 0Gy_TLR3i_10uM vs. 0Gy_TLR4i_10uM | -0.6035 | -11.80 to 10.59 | ns | >0.9999 |
| 0Gy_TLR3i_10uM vs. 0Gy_CQ_1uM | 0.4179 | -10.78 to 11.61 | ns | >0.9999 |
| 0Gy_TLR3i_10uM vs. 0Gy_CQ_5uM | 0.7977 | -10.40 to 11.99 | ns | >0.9999 |
| 0Gy_TLR3i_10uM vs. 0Gy_CQ_10uM | 0.6101 | -10.59 to 11.81 | ns | >0.9999 |
| 0Gy_TLR3i_10uM vs. 4Gy_DMSO | -13.80 | -25.00 to -2.608 | ** | 0.0045 |
| 0Gy_TLR3i_10uM vs. 4Gy_TLR3i_5uM | -6.967 | -18.16 to 4.229 | ns | 0.6606 |
| 0Gy_TLR3i_10uM vs. 4Gy_TLR3i_10uM | -2.545 | -13.74 to 8.650 | ns | >0.9999 |
| 0Gy_TLR3i_10uM vs. 4Gy_TLR4i_5uM | -15.91 | -27.11 to -4.715 | *** | 0.0005 |
| 0Gy_TLR3i_10uM vs. 4Gy_TLR4i_10uM | -9.482 | -20.68 to 1.714 | ns | 0.1836 |
| 0Gy_TLR3i_10uM vs. 4Gy_CQ_1uM | -0.01571 | -11.21 to 11.18 | ns | >0.9999 |
| 0Gy_TLR3i_10uM vs. 4Gy_CQ_5uM | 0.9732 | -10.22 to 12.17 | ns | >0.9999 |
| 0Gy_TLR3i_10uM vs. 4Gy_CQ_10uM | 0.9728 | -10.22 to 12.17 | ns | >0.9999 |
| 0Gy_TLR4i_5uM vs. 0Gy_TLR4i_10uM | -0.9065 | -12.10 to 10.29 | ns | >0.9999 |
| 0Gy_TLR4i_5uM vs. 0Gy_CQ_1uM | 0.1150 | -11.08 to 11.31 | ns | >0.9999 |
| 0Gy_TLR4i_5uM vs. 0Gy_CQ_5uM | 0.4948 | -10.70 to 11.69 | ns | >0.9999 |
| 0Gy_TLR4i_5uM vs. 0Gy_CQ_10uM | 0.3072 | -10.89 to 11.50 | ns | >0.9999 |
| 0Gy_TLR4i_5uM vs. 4Gy_DMSO | -14.11 | -25.30 to -2.911 | ** | 0.0033 |
| 0Gy_TLR4i_5uM vs. 4Gy_TLR3i_5uM | -7.270 | -18.47 to 3.926 | ns | 0.5939 |
| 0Gy_TLR4i_5uM vs. 4Gy_TLR3i_10uM | -2.848 | -14.04 to 8.348 | ns | 0.9999 |
| 0Gy_TLR4i_5uM vs. 4Gy_TLR4i_5uM | -16.21 | -27.41 to -5.018 | *** | 0.0004 |
| 0Gy_TLR4i_5uM vs. 4Gy_TLR4i_10uM | -9.785 | -20.98 to 1.411 | ns | 0.1490 |

| | | | | |
|---|---|---|---|---|
| 0Gy_TLR4i_5uM vs. 4Gy_CQ_1uM | -0.3186 | -11.51 to 10.88 | ns | >0.9999 |
| 0Gy_TLR4i_5uM vs. 4Gy_CQ_5uM | 0.6703 | -10.53 to 11.87 | ns | >0.9999 |
| 0Gy_TLR4i_5uM vs. 4Gy_CQ_10uM | 0.6698 | -10.53 to 11.87 | ns | >0.9999 |
| 0Gy_TLR4i_10uM vs. 0Gy_CQ_1uM | 1.021 | -10.17 to 12.22 | ns | >0.9999 |
| 0Gy_TLR4i_10uM vs. 0Gy_CQ_5uM | 1.401 | -9.795 to 12.60 | ns | >0.9999 |
| 0Gy_TLR4i_10uM vs. 0Gy_CQ_10uM | 1.214 | -9.982 to 12.41 | ns | >0.9999 |
| 0Gy_TLR4i_10uM vs. 4Gy_DMSO | -13.20 | -24.40 to -2.005 | ** | 0.0081 |
| 0Gy_TLR4i_10uM vs. 4Gy_TLR3i_5uM | -6.364 | -17.56 to 4.832 | ns | 0.7835 |
| 0Gy_TLR4i_10uM vs. 4Gy_TLR3i_10uM | -1.942 | -13.14 to 9.254 | ns | >0.9999 |
| 0Gy_TLR4i_10uM vs. 4Gy_TLR4i_5uM | -15.31 | -26.50 to -4.112 | *** | 0.0010 |
| 0Gy_TLR4i_10uM vs. 4Gy_TLR4i_10uM | -8.879 | -20.07 to 2.317 | ns | 0.2696 |
| 0Gy_TLR4i_10uM vs. 4Gy_CQ_1uM | 0.5878 | -10.61 to 11.78 | ns | >0.9999 |
| 0Gy_TLR4i_10uM vs. 4Gy_CQ_5uM | 1.577 | -9.619 to 12.77 | ns | >0.9999 |
| 0Gy_TLR4i_10uM vs. 4Gy_CQ_10uM | 1.576 | -9.620 to 12.77 | ns | >0.9999 |
| 0Gy_CQ_1uM vs. 0Gy_CQ_5uM | 0.3797 | -10.82 to 11.58 | ns | >0.9999 |
| 0Gy_CQ_1uM vs. 0Gy_CQ_10uM | 0.1922 | -11.00 to 11.39 | ns | >0.9999 |
| 0Gy_CQ_1uM vs. 4Gy_DMSO | -14.22 | -25.42 to -3.026 | ** | 0.0030 |
| 0Gy_CQ_1uM vs. 4Gy_TLR3i_5uM | -7.385 | -18.58 to 3.811 | ns | 0.5683 |
| 0Gy_CQ_1uM vs. 4Gy_TLR3i_10uM | -2.963 | -14.16 to 8.233 | ns | 0.9998 |
| 0Gy_CQ_1uM vs. 4Gy_TLR4i_5uM | -16.33 | -27.53 to -5.133 | *** | 0.0003 |
| 0Gy_CQ_1uM vs. 4Gy_TLR4i_10uM | -9.900 | -21.10 to 1.296 | ns | 0.1373 |
| 0Gy_CQ_1uM vs. 4Gy_CQ_1uM | -0.4337 | -11.63 to 10.76 | ns | >0.9999 |
| 0Gy_CQ_1uM vs. 4Gy_CQ_5uM | 0.5553 | -10.64 to 11.75 | ns | >0.9999 |
| 0Gy_CQ_1uM vs. 4Gy_CQ_10uM | 0.5548 | -10.64 to 11.75 | ns | >0.9999 |
| 0Gy_CQ_5uM vs. 0Gy_CQ_10uM | -0.1876 | -11.38 to 11.01 | ns | >0.9999 |
| 0Gy_CQ_5uM vs. 4Gy_DMSO | -14.60 | -25.80 to -3.406 | ** | 0.0020 |
| 0Gy_CQ_5uM vs. 4Gy_TLR3i_5uM | -7.765 | -18.96 to 3.431 | ns | 0.4845 |

| | | | | |
|---|---|---|---|---|
| 0Gy_CQ_5uM vs. 4Gy_TLR3i_10uM | -3.343 | -14.54 to 7.853 | ns | 0.9992 |
| 0Gy_CQ_5uM vs. 4Gy_TLR4i_5uM | -16.71 | -27.90 to -5.513 | *** | 0.0002 |
| 0Gy_CQ_5uM vs. 4Gy_TLR4i_10uM | -10.28 | -21.48 to 0.9160 | ns | 0.1038 |
| 0Gy_CQ_5uM vs. 4Gy_CQ_1uM | -0.8134 | -12.01 to 10.38 | ns | >0.9999 |
| 0Gy_CQ_5uM vs. 4Gy_CQ_5uM | 0.1755 | -11.02 to 11.37 | ns | >0.9999 |
| 0Gy_CQ_5uM vs. 4Gy_CQ_10uM | 0.1751 | -11.02 to 11.37 | ns | >0.9999 |
| 0Gy_CQ_10uM vs. 4Gy_DMSO | -14.41 | -25.61 to -3.218 | ** | 0.0024 |
| 0Gy_CQ_10uM vs. 4Gy_TLR3i_5uM | -7.577 | -18.77 to 3.619 | ns | 0.5256 |
| 0Gy_CQ_10uM vs. 4Gy_TLR3i_10uM | -3.156 | -14.35 to 8.040 | ns | 0.9996 |
| 0Gy_CQ_10uM vs. 4Gy_TLR4i_5uM | -16.52 | -27.72 to -5.325 | *** | 0.0003 |
| 0Gy_CQ_10uM vs. 4Gy_TLR4i_10uM | -10.09 | -21.29 to 1.104 | ns | 0.1194 |
| 0Gy_CQ_10uM vs. 4Gy_CQ_1uM | -0.6258 | -11.82 to 10.57 | ns | >0.9999 |
| 0Gy_CQ_10uM vs. 4Gy_CQ_5uM | 0.3631 | -10.83 to 11.56 | ns | >0.9999 |
| 0Gy_CQ_10uM vs. 4Gy_CQ_10uM | 0.3627 | -10.83 to 11.56 | ns | >0.9999 |
| 4Gy_DMSO vs. 4Gy_TLR3i_5uM | 6.837 | -4.359 to 18.03 | ns | 0.6886 |
| 4Gy_DMSO vs. 4Gy_TLR3i_10uM | 11.26 | 0.06279 to 22.45 | * | 0.0474 |
| 4Gy_DMSO vs. 4Gy_TLR4i_5uM | -2.107 | -13.30 to 9.089 | ns | >0.9999 |
| 4Gy_DMSO vs. 4Gy_TLR4i_10uM | 4.322 | -6.874 to 15.52 | ns | 0.9882 |
| 4Gy_DMSO vs. 4Gy_CQ_1uM | 13.79 | 2.593 to 24.98 | ** | 0.0046 |
| 4Gy_DMSO vs. 4Gy_CQ_5uM | 14.78 | 3.581 to 25.97 | ** | 0.0017 |
| 4Gy_DMSO vs. 4Gy_CQ_10uM | 14.78 | 3.581 to 25.97 | ** | 0.0017 |
| 4Gy_TLR3i_5uM vs. 4Gy_TLR3i_10uM | 4.422 | -6.774 to 15.62 | ns | 0.9854 |
| 4Gy_TLR3i_5uM vs. 4Gy_TLR4i_5uM | -8.944 | -20.14 to 2.252 | ns | 0.2592 |
| 4Gy_TLR3i_5uM vs. 4Gy_TLR4i_10uM | -2.515 | -13.71 to 8.681 | ns | >0.9999 |
| 4Gy_TLR3i_5uM vs. 4Gy_CQ_1uM | 6.952 | -4.244 to 18.15 | ns | 0.6640 |
| 4Gy_TLR3i_5uM vs. 4Gy_CQ_5uM | 7.940 | -3.255 to 19.14 | ns | 0.4468 |
| 4Gy_TLR3i_5uM vs. 4Gy_CQ_10uM | 7.940 | -3.256 to 19.14 | ns | 0.4469 |

| | | | | |
|---|---|---|---|---|
| 4Gy_TLR3i_10uM vs. 4Gy_TLR4i_5uM | -13.37 | -24.56 to -2.170 | ** | 0.0069 |
| 4Gy_TLR3i_10uM vs. 4Gy_TLR4i_10uM | -6.937 | -18.13 to 4.259 | ns | 0.6672 |
| 4Gy_TLR3i_10uM vs. 4Gy_CQ_1uM | 2.530 | -8.666 to 13.73 | ns | >0.9999 |
| 4Gy_TLR3i_10uM vs. 4Gy_CQ_5uM | 3.519 | -7.677 to 14.71 | ns | 0.9986 |
| 4Gy_TLR3i_10uM vs. 4Gy_CQ_10uM | 3.518 | -7.678 to 14.71 | ns | 0.9986 |
| 4Gy_TLR4i_5uM vs. 4Gy_TLR4i_10uM | 6.429 | -4.767 to 17.62 | ns | 0.7712 |
| 4Gy_TLR4i_5uM vs. 4Gy_CQ_1uM | 15.90 | 4.700 to 27.09 | *** | 0.0005 |
| 4Gy_TLR4i_5uM vs. 4Gy_CQ_5uM | 16.88 | 5.688 to 28.08 | *** | 0.0002 |
| 4Gy_TLR4i_5uM vs. 4Gy_CQ_10uM | 16.88 | 5.688 to 28.08 | *** | 0.0002 |
| 4Gy_TLR4i_10uM vs. 4Gy_CQ_1uM | 9.467 | -1.729 to 20.66 | ns | 0.1855 |
| 4Gy_TLR4i_10uM vs. 4Gy_CQ_5uM | 10.46 | -0.7405 to 21.65 | ns | 0.0908 |
| 4Gy_TLR4i_10uM vs. 4Gy_CQ_10uM | 10.45 | -0.7410 to 21.65 | ns | 0.0908 |
| 4Gy_CQ_1uM vs. 4Gy_CQ_5uM | 0.9889 | -10.21 to 12.18 | ns | >0.9999 |
| 4Gy_CQ_1uM vs. 4Gy_CQ_10uM | 0.9885 | -10.21 to 12.18 | ns | >0.9999 |
| 4Gy_CQ_5uM vs. 4Gy_CQ_10uM | -0.0004362 | -11.20 to 11.20 | ns | >0.9999 |

**Supplementary Table 8.** Two-way ANOVA analysis results of multiple comparisons for HK-374 data represented in Figure 4B (RT-PCR for YFs following IR+TLR3/4i).

| Tukey's multiple comparisons test | Predicted (LS) mean diff. | 95.00% CI of diff. | Summary | Adjusted P Value |
|---|---|---|---|---|
| **0ct4** | | | | |
| 0Gy_DMSO vs. 4Gy_DMSO | -0.9651 | -2.768 to 0.8382 | ns | 0.5061 |
| 0Gy_DMSO vs. 4Gy_TLR3i_5uM | -0.02787 | -1.674 to 1.618 | ns | >0.9999 |
| 0Gy_DMSO vs. 4Gy_TLR4i_5uM | -0.1613 | -1.808 to 1.485 | ns | 0.9942 |
| 4Gy_DMSO vs. 4Gy_TLR3i_5uM | 0.9373 | -0.7089 to 2.583 | ns | 0.4514 |
| 4Gy_DMSO vs. 4Gy_TLR4i_5uM | 0.8038 | -0.8424 to 2.450 | ns | 0.5831 |
| 4Gy_TLR3i_5uM vs. 4Gy_TLR4i_5uM | -0.1334 | -1.606 to 1.339 | ns | 0.9954 |
| **Sox2** | | | | |
| 0Gy_DMSO vs. 4Gy_DMSO | -0.4973 | -2.301 to 1.306 | ns | 0.8899 |
| 0Gy_DMSO vs. 4Gy_TLR3i_5uM | 0.07439 | -1.572 to 1.721 | ns | 0.9994 |
| 0Gy_DMSO vs. 4Gy_TLR4i_5uM | 0.1882 | -1.458 to 1.834 | ns | 0.9908 |
| 4Gy_DMSO vs. 4Gy_TLR3i_5uM | 0.5717 | -1.074 to 2.218 | ns | 0.8028 |
| 4Gy_DMSO vs. 4Gy_TLR4i_5uM | 0.6855 | -0.9607 to 2.332 | ns | 0.7001 |
| 4Gy_TLR3i_5uM vs. 4Gy_TLR4i_5uM | 0.1138 | -1.359 to 1.586 | ns | 0.9971 |
| **Klf4** | | | | |
| 0Gy_DMSO vs. 4Gy_DMSO | -2.507 | -4.310 to -0.7035 | ** | 0.0024 |
| 0Gy_DMSO vs. 4Gy_TLR3i_5uM | -1.061 | -2.708 to 0.5848 | ns | 0.3394 |
| 0Gy_DMSO vs. 4Gy_TLR4i_5uM | -0.9295 | -2.576 to 0.7167 | ns | 0.4589 |
| 4Gy_DMSO vs. 4Gy_TLR3i_5uM | 1.445 | -0.2008 to 3.092 | ns | 0.1068 |
| 4Gy_DMSO vs. 4Gy_TLR4i_5uM | 1.577 | -0.06888 to 3.224 | ns | 0.0656 |
| 4Gy_TLR3i_5uM vs. 4Gy_TLR4i_5uM | 0.1319 | -1.340 to 1.604 | ns | 0.9955 |
| **Nanog** | | | | |
| 0Gy_DMSO vs. 4Gy_DMSO | -3.336 | -5.139 to -1.533 | **** | <0.0001 |
| 0Gy_DMSO vs. 4Gy_TLR3i_5uM | -0.1864 | -1.833 to 1.460 | ns | 0.9911 |

| | | | | |
|---|---|---|---|---|
| 0Gy_DMSO vs. 4Gy_TLR4i_5uM | -0.4116 | -2.058 to 1.235 | ns | 0.9151 |
| 4Gy_DMSO vs. 4Gy_TLR3i_5uM | 3.150 | 1.503 to 4.796 | **** | <0.0001 |
| 4Gy_DMSO vs. 4Gy_TLR4i_5uM | 2.924 | 1.278 to 4.571 | **** | <0.0001 |
| 4Gy_TLR3i_5uM vs. 4Gy_TLR4i_5uM | -0.2252 | -1.698 to 1.247 | ns | 0.9786 |
| **cMyc** | | | | |
| 0Gy_DMSO vs. 4Gy_DMSO | -1.882 | -3.685 to -0.07849 | * | 0.0372 |
| 0Gy_DMSO vs. 4Gy_TLR3i_5uM | -0.6269 | -2.273 to 1.019 | ns | 0.7547 |
| 0Gy_DMSO vs. 4Gy_TLR4i_5uM | -0.5266 | -2.173 to 1.120 | ns | 0.8389 |
| 4Gy_DMSO vs. 4Gy_TLR3i_5uM | 1.255 | -0.3913 to 2.901 | ns | 0.1994 |
| 4Gy_DMSO vs. 4Gy_TLR4i_5uM | 1.355 | -0.2910 to 3.001 | ns | 0.1453 |
| 4Gy_TLR3i_5uM vs. 4Gy_TLR4i_5uM | 0.1003 | -1.372 to 1.573 | ns | 0.9980 |

**Supplementary Table 9.** Two-way ANOVA analysis results of multiple comparisons for data represented in Figure 5A (ELDA for cGASi).

| Dunnett's multiple comparisons test | Mean Diff. | 95.00% CI of diff. | Summary | Adjusted P Value |
|---|---|---|---|---|
| **0Gy** | | | | |
| DMSO vs. G140 5uM | -0.1047 | -0.3269 to 0.1176 | ns | 0.4464 |
| DMSO vs. G140 10uM | -0.06260 | -0.2849 to 0.1597 | ns | 0.7354 |
| **2Gy** | | | | |
| DMSO vs. G140 5uM | -0.09900 | -0.3213 to 0.1233 | ns | 0.4828 |
| DMSO vs. G140 10uM | -0.01533 | -0.2376 to 0.2069 | ns | 0.9810 |
| **4Gy** | | | | |
| DMSO vs. G140 5uM | -0.001352 | -0.2236 to 0.2209 | ns | 0.9999 |
| DMSO vs. G140 10uM | 0.06053 | -0.1617 to 0.2828 | ns | 0.7495 |
| **8Gy** | | | | |
| DMSO vs. G140 5uM | 0.02182 | -0.2004 to 0.2441 | ns | 0.9619 |
| DMSO vs. G140 10uM | 0.003459 | -0.2188 to 0.2257 | ns | 0.9990 |

**Supplementary Table 10.** Two-way ANOVA analysis results of multiple comparisons for HK-374 glioma sphere data represented in Figure 5B (ELDA for STINGi).

| Dunnett's multiple comparisons test | Predicted (LS) mean diff. | 95.00% CI of diff. | Summary | Adjusted P Value |
|---|---|---|---|---|
| **0Gy** | | | | |
| DMSO vs. H-151 250nM | 0.2400 | 0.06115 to 0.4188 | ** | 0.0057 |
| DMSO vs. H-151 500nM | 0.4465 | 0.2677 to 0.6254 | **** | <0.0001 |
| DMSO vs. H-151 1uM | 0.8710 | 0.6922 to 1.050 | **** | <0.0001 |
| **2Gy** | | | | |
| DMSO vs. H-151 250nM | 0.3203 | 0.08943 to 0.5512 | ** | 0.0041 |
| DMSO vs. H-151 500nM | 0.4860 | 0.2551 to 0.7169 | **** | <0.0001 |
| DMSO vs. H-151 1uM | 0.6491 | 0.4182 to 0.8800 | **** | <0.0001 |
| **4Gy** | | | | |
| DMSO vs. H-151 250nM | 0.1279 | -0.05098 to 0.3067 | ns | 0.2128 |
| DMSO vs. H-151 500nM | 0.1672 | -0.01164 to 0.3460 | ns | 0.0719 |
| DMSO vs. H-151 1uM | 0.2148 | 0.03595 to 0.3936 | * | 0.0146 |
| **8Gy** | | | | |
| DMSO vs. H-151 250nM | 0.008023 | -0.1708 to 0.1869 | ns | 0.9991 |
| DMSO vs. H-151 500nM | 0.01169 | -0.1672 to 0.1905 | ns | 0.9972 |
| DMSO vs. H-151 1uM | 0.01672 | -0.1621 to 0.1956 | ns | 0.9920 |

**Supplementary Table 11.** Two-way ANOVA analysis results of multiple comparisons for HK-157 glioma sphere data represented in Figure 5C (ELDA for STINGi).

| Dunnett's multiple comparisons test | Mean Diff. | 95.00% CI of diff. | Summary | Adjusted P Value |
|---|---|---|---|---|
| **0Gy** | | | | |
| DMSO vs. H-151 250nM | 0.2902 | 0.1378 to 0.4426 | *** | 0.0001 |
| DMSO vs. H-151 500nM | 0.3141 | 0.1617 to 0.4665 | **** | <0.0001 |
| DMSO vs. H-151 1uM | 0.7285 | 0.5761 to 0.8809 | **** | <0.0001 |
| **2Gy** | | | | |
| DMSO vs. H-151 250nM | 0.07060 | -0.08179 to 0.2230 | ns | 0.5334 |
| DMSO vs. H-151 500nM | 0.1811 | 0.02873 to 0.3335 | * | 0.0168 |
| DMSO vs. H-151 1uM | 0.2611 | 0.1087 to 0.4135 | *** | 0.0006 |
| **4Gy** | | | | |
| DMSO vs. H-151 250nM | 0.02204 | -0.1304 to 0.1744 | ns | 0.9700 |
| DMSO vs. H-151 500nM | 0.04105 | -0.1113 to 0.1934 | ns | 0.8455 |
| DMSO vs. H-151 1uM | 0.07677 | -0.07563 to 0.2292 | ns | 0.4688 |
| **8Gy** | | | | |
| DMSO vs. H-151 250nM | -0.007944 | -0.1603 to 0.1444 | ns | 0.9985 |
| DMSO vs. H-151 500nM | 0.002831 | -0.1496 to 0.1552 | ns | >0.9999 |
| DMSO vs. H-151 1uM | 0.002830 | -0.1496 to 0.1552 | ns | >0.9999 |

**Supplementary Table 12.** Two-way ANOVA analysis results of multiple comparisons for HK-308 glioma sphere data represented in Figure 5D (ELDA for STINGi).

| Dunnett's multiple comparisons test | Mean Diff. | 95.00% CI of diff. | Summary | Adjusted P Value |
|---|---|---|---|---|
| **0Gy** | | | | |
| DMSO vs. H-151 250nM | 0.2823 | 0.1404 to 0.4242 | **** | <0.0001 |
| DMSO vs. H-151 500nM | 0.1871 | 0.04519 to 0.3291 | ** | 0.0075 |
| DMSO vs. H-151 1uM | 0.4791 | 0.3371 to 0.6210 | **** | <0.0001 |
| **2Gy** | | | | |
| DMSO vs. H-151 250nM | 0.02493 | -0.1170 to 0.1669 | ns | 0.9485 |
| DMSO vs. H-151 500nM | 0.004924 | -0.1370 to 0.1469 | ns | 0.9995 |
| DMSO vs. H-151 1uM | 0.05644 | -0.08549 to 0.1984 | ns | 0.6429 |
| **4Gy** | | | | |
| DMSO vs. H-151 250nM | -0.005924 | -0.1479 to 0.1360 | ns | 0.9992 |
| DMSO vs. H-151 500nM | 0.005658 | -0.1363 to 0.1476 | ns | 0.9993 |
| DMSO vs. H-151 1uM | 0.03063 | -0.1113 to 0.1726 | ns | 0.9111 |
| **8Gy** | | | | |
| DMSO vs. H-151 250nM | -0.01104 | -0.1530 to 0.1309 | ns | 0.9950 |
| DMSO vs. H-151 500nM | 0.003566 | -0.1384 to 0.1455 | ns | 0.9999 |
| DMSO vs. H-151 1uM | -0.001506 | -0.1434 to 0.1404 | ns | >0.9999 |

**Supplementary Table 13.** Two-way ANOVA analysis results of multiple comparisons for HK-374 glioma sphere data represented in Figure 5E (SC freq for STINGi ELDAs).

| Dunnett's multiple comparisons test | Predicted (LS) mean diff. | 95.00% CI of diff. | Summary | Adjusted P Value |
|---|---|---|---|---|
| **0Gy** | | | | |
| DMSO vs. new H-151 250nm | 12.19 | 1.878 to 22.51 | * | 0.0165 |
| DMSO vs. new H-151 500nm | 31.77 | 21.46 to 42.09 | **** | <0.0001 |
| DMSO vs. new H-151 1uM | 45.13 | 34.82 to 55.45 | **** | <0.0001 |
| **2Gy** | | | | |
| DMSO vs. new H-151 250nm | 12.42 | -0.8966 to 25.74 | ns | 0.0727 |
| DMSO vs. new H-151 500nm | 19.41 | 6.093 to 32.73 | ** | 0.0025 |
| DMSO vs. new H-151 1uM | 21.46 | 8.146 to 34.78 | *** | 0.0008 |
| **4Gy** | | | | |
| DMSO vs. new H-151 250nm | 3.705 | -6.610 to 14.02 | ns | 0.7186 |
| DMSO vs. new H-151 500nm | 5.634 | -4.681 to 15.95 | ns | 0.4165 |
| DMSO vs. new H-151 1uM | 6.423 | -3.892 to 16.74 | ns | 0.3130 |
| **8Gy** | | | | |
| DMSO vs. new H-151 250nm | 0.1582 | -10.16 to 10.47 | ns | >0.9999 |
| DMSO vs. new H-151 500nm | 0.4771 | -9.838 to 10.79 | ns | 0.9990 |
| DMSO vs. new H-151 1uM | 0.5766 | -9.738 to 10.89 | ns | 0.9982 |

**Supplementary Table 14.** Two-way ANOVA analysis results of multiple comparisons for HK-157 glioma sphere data represented in Figure 5F (SC freq for STINGi ELDAs).

| Dunnett's multiple comparisons test | Mean Diff. | 95.00% CI of diff. | Summary | Adjusted P Value |
|---|---|---|---|---|
| **0Gy** | | | | |
| DMSO vs. new H-151 250nm | 1.714 | -3.716 to 7.144 | ns | 0.7775 |
| DMSO vs. new H-151 500nm | 11.55 | 6.124 to 16.98 | **** | <0.0001 |
| DMSO vs. new H-151 1uM | 15.02 | 9.591 to 20.45 | **** | <0.0001 |
| **2Gy** | | | | |
| DMSO vs. new H-151 250nm | 0.6832 | -4.747 to 6.113 | ns | 0.9797 |
| DMSO vs. new H-151 500nm | 1.754 | -3.676 to 7.184 | ns | 0.7659 |
| DMSO vs. new H-151 1uM | 2.517 | -2.913 to 7.947 | ns | 0.5330 |
| **4Gy** | | | | |
| DMSO vs. new H-151 250nm | -0.1392 | -5.569 to 5.291 | ns | 0.9999 |
| DMSO vs. new H-151 500nm | 0.5467 | -4.883 to 5.977 | ns | 0.9894 |
| DMSO vs. new H-151 1uM | 0.7925 | -4.638 to 6.223 | ns | 0.9692 |
| **8Gy** | | | | |
| DMSO vs. new H-151 250nm | -0.02856 | -5.459 to 5.402 | ns | >0.9999 |
| DMSO vs. new H-151 500nm | 0.02078 | -5.409 to 5.451 | ns | >0.9999 |
| DMSO vs. new H-151 1uM | 0.01684 | -5.413 to 5.447 | ns | >0.9999 |

**Supplementary Table 15.** Two-way ANOVA analysis results of multiple comparisons for HK-308 glioma sphere data represented in Figure 5G(SC freq for STINGi ELDAs).

| Dunnett's multiple comparisons test | Mean Diff. | 95.00% CI of diff. | Summary | Adjusted P Value |
|---|---|---|---|---|
| **0Gy** | | | | |
| DMSO vs. new H-151 250nm | 3.980 | -1.295 to 9.255 | ns | 0.1729 |
| DMSO vs. new H-151 500nm | 8.174 | 2.899 to 13.45 | ** | 0.0016 |
| DMSO vs. new H-151 1uM | 9.838 | 4.563 to 15.11 | *** | 0.0002 |
| **2Gy** | | | | |
| DMSO vs. new H-151 250nm | 0.02501 | -5.250 to 5.300 | ns | >0.9999 |
| DMSO vs. new H-151 500nm | -0.2250 | -5.500 to 5.050 | ns | 0.9991 |
| DMSO vs. new H-151 1uM | 0.6788 | -4.596 to 5.954 | ns | 0.9784 |
| **4Gy** | | | | |
| DMSO vs. new H-151 250nm | -0.3726 | -5.648 to 4.902 | ns | 0.9962 |
| DMSO vs. new H-151 500nm | -0.1475 | -5.422 to 5.127 | ns | 0.9997 |
| DMSO vs. new H-151 1uM | 0.01350 | -5.261 to 5.288 | ns | >0.9999 |
| **8Gy** | | | | |
| DMSO vs. new H-151 250nm | -0.04609 | -5.321 to 5.229 | ns | >0.9999 |
| DMSO vs. new H-151 500nm | 0.007596 | -5.267 to 5.283 | ns | >0.9999 |
| DMSO vs. new H-151 1uM | -0.02876 | -5.304 to 5.246 | ns | >0.9999 |
